# Quantitative real-time in-cell imaging reveals heterogeneous clusters of proteins prior to condensation

**DOI:** 10.1101/2022.08.01.502196

**Authors:** Chenyang Lan, Juhyeong Kim, Svenja Ulferts, Fernando Aprile-Garcia, Abhinaya Anandamurugan, Robert Grosse, Ritwick Sawarkar, Aleks Reinhardt, Thorsten Hugel

## Abstract

The formation of biomolecular condensates underpins many cellular processes; however, our current understanding of condensate formation within cells is largely based on observing the final near-equilibrium condensate state. It is less clear how proteins behave before condensates form or at concentrations at which condensation does not occur in cells. Here, we use a combination of fluorescence microscopy and photobleaching analysis to quantify phase separation of negative elongation factor (NELF) in living and stressed cells. We use the recently reported system of stress-induced condensation of NELF in human nuclei as a model to study the behaviour of proteins before condensation. We find that pre-condensate heterogeneous clusters both grow and shrink and are not freely diffusing. Unexpectedly, we also find such small dynamic clusters in unstressed cells in which condensates do not form. We provide a categorisation of small and large clusters based on their dynamics and their response to p38 kinase inhibition. Overall, our data are best explained as non-classical nucleation with a flat free-energy landscape for clusters of a range of sizes and an inhibition of condensation.

---

Recent work has shown that numerous cellular processes are underpinned by the formation of biomolecular condensates [1, 2]. Key steps of gene expression, including transcription [3–6], translation, as well as signalling [7–10] and metabolism [11, 12], are regulated by membraneless assemblies of relevant macromolecules, namely proteins and nucleic acids. Such membraneless assemblies have the advantage of rapid material exchange with their surroundings while keeping the macromolecules in spatial proximity [13, 14]. They are thought to form by phase separation where the macromolecular concentration is much higher within them than in their immediate surroundings [15, 16], and are therefore often called condensates. Intriguingly, misregulated condensation has been shown to be causally linked with human pathologies [16–22]. A molecular understanding of the process of condensation is thus important from both a fundamental [16, 23–27] and a biomedical perspective [28, 29].

A key question in the field is how proteins form condensates at a molecular level [30, 31]. This question can be broken down into two related issues. First, it is not clear how proteins behave prior to the formation of condensates or at subsaturated concentrations at which condensates do not form. Second, it would be useful to know what properties of proteins change during condensate formation. In the traditional homogeneous nucleation picture of condensate formation, there is a competition between a favourable bulk term and a disfavourable surface term associated with the formation of an interface between the cluster and its surroundings [32]. The free energy of a cluster rapidly increases until a critical cluster size is reached and the bulk term begins to dominate [32]; clusters are therefore most likely either post-critical or very small [30]. In subsaturated conditions, the bulk term is itself disfavourable and only small fluctuations in cluster size around the homogeneous monomeric phase occur. Protein behaviour in such conditions is actively being investigated *in vitro* using a fixed uniform concentration of recombinant pure components, and direct observation of the protein dynamics in purified systems allows for building and testing theoretical models [33]. Interestingly, small clusters of a range of sizes were observed under subsaturation conditions, suggesting that a simple two-state description of the system is insufficient. If similar long-lived small clusters of more than a few molecules are also present in cells, this may fundamentally change our understanding of the physical basis of biological phase separation. However, condensate formation within the complex cellular milieu of diverse macromolecules with variable concentrations has been difficult to address. Molecular crowding, the existence of inflexible polymers such as the cytoskeleton and a plethora of small-molecule metabolites limit the extrapolation of *in vitro* studies to cellular condensates. Moreover, the non-equilibrium environment of the cell allows for circumvention of thermodynamic constraints and the emergence of new features, such as dynamic droplet localization, which can arise in active systems [34].

Despite some recent successes [1, 4, 5, 35–39], the quantification of the dynamics of phase separation in living cells is still difficult, perhaps because the study of proteins prior to condensate formation inside living cells is limited by several technical impediments. First, the signal-to-noise ratio in fluorescence measurements in living cells is low, and counting the number of fluorescent proteins is therefore usually done in fixed cells. Second, the density of proteins in clusters is high, which impedes the separation and counting of single proteins even with super-resolution imaging [4]. Third, proteins within cells are mobile and dynamic, necessitating a high time resolution that is currently difficult to achieve with commercial setups at a low signal-to-noise ratio. In addition to these technical reasons, it is difficult to capture proteins in their non-equilibrium ‘pre-condensate state’. While *in vitro* studies can rely on titrating concentrations of proteins below the saturation threshold to observe such states, it is not straightforward to control protein levels inside cells. Given these limitations, most studies thus far have largely focused on the late (‘equilibrium’) states of condensates even when small transient clusters or oligomers prior to condensate formation were detected [4, 40], leaving a gap in our understanding of the pre-condensate behaviour of proteins.

The observation of pre-condensate protein behaviour within cells requires a controlled, signalling-induced transition of proteins to condensates, separating condensate and pre-condensate states in time. An example of such a process is the stress-induced condensation of a nuclear transcriptional regulator, negative elongation factor (NELF). NELF has been closely linked to stress-induced transcriptional attenuation (SITA); moreover, p38 kinase signalling has been connected to gene downregulation [41]. Simple and controlled stressors such as As_2_O_3_ cause NELF to form condensates, leading to a global downregulation of transcription [42]. The NELF complex comprises four subunits, with NELFA possessing an intrinsically disordered region and NELFE a receptor-binding domain [41, 43]. Expression of NELFA-GFP enables NELF condensation to be observed upon arsenic stress in real time in living cells. In this study, we combine super-resolution imaging and single-molecule microscopy in fixed and living cells to quantify the behaviour of NELF in cells both before and during condensation. We also investigate the effect of a p38 kinase inhibitor [41] on this process.

## RESULTS AND DISCUSSION

### Dynamic and transient clusters can be tracked and quantified in living cells

Tracking cluster growth at near single-molecule sensitivity and a high time resolution requires the overall concentration of the tracked molecules to be low. To this end, we used a tetracycline-inducible system in HeLa cells to achieve low expression levels of NELFA-GFP [42]. We identified conditions under which NELFA-GFP is expressed to levels of ~25 % of the endogenous NELFA in HeLa cells (Suppl. Note 1). The condensation of NELFA was triggered by toxic stress (100 μM As_2_O_3_), which has been shown to result in similar condensation as heat stress [42]. We ensured that this treatment did not compromise cell viability (Suppl. Note 2).

To test the effect of NELFA-GFP concentration on condensation, we first imaged cell nuclei at high and low expression conditions before and after exposure to stress [Fig. 1]. At low expression conditions, upon exposure to arsenic, several small clusters of NELFA-GFP, but only a single large condensate, were visible, while many large condensates occurred at high expression conditions. Such a dependence on concentration is expected for condensate formation and therefore in the following we work at the lowest expression conditions, i.e. where a single large condensate forms, in order to be able to track single small clusters.

**Figure 1.**
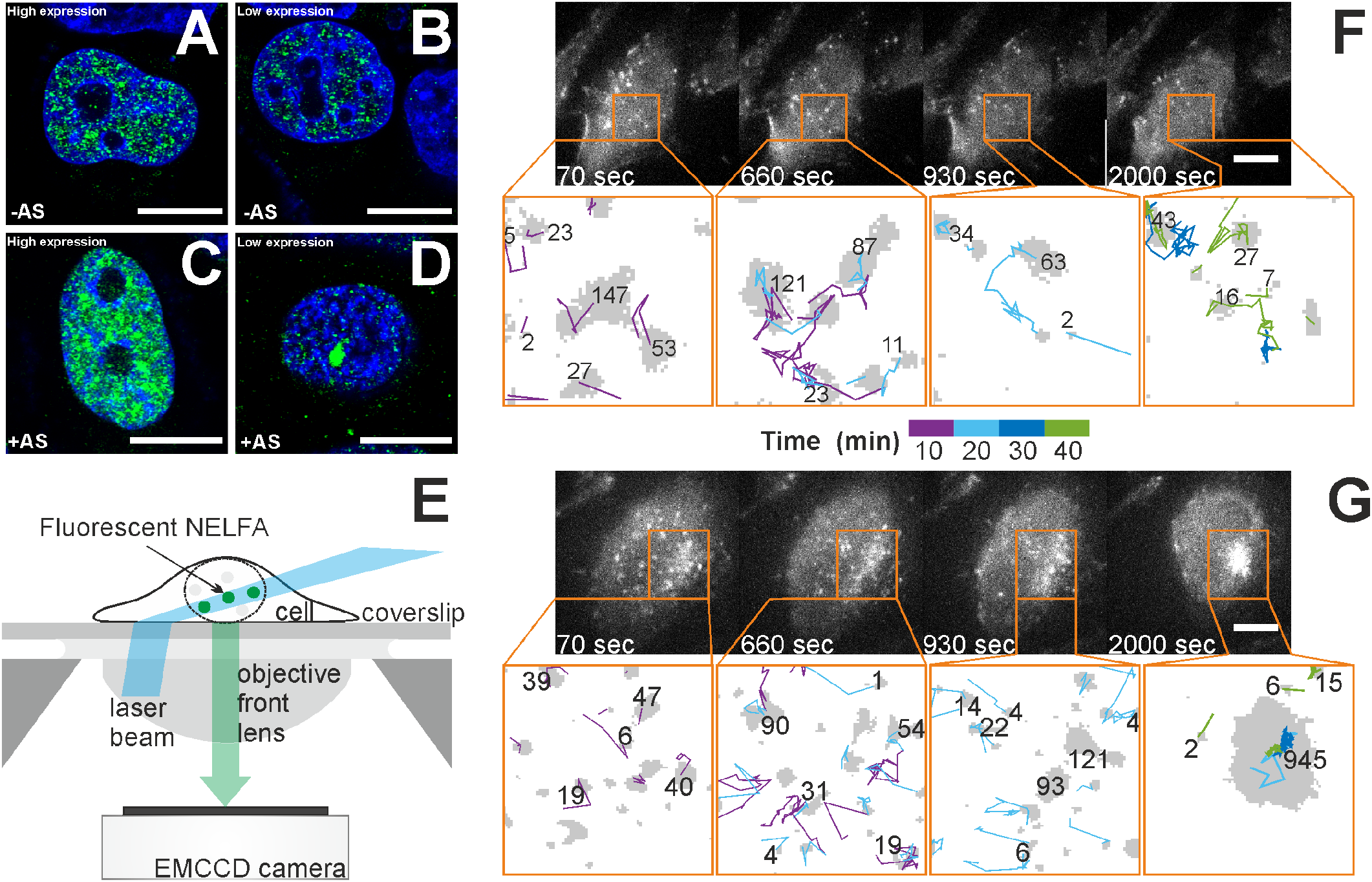
Time-resolved imaging of clusters and condensates in cells. **A–D** High-resolution Airyscan 3D-imaging of HeLa cells with NELFA-GFP at standard (**A** and **C**) and low (**B** and **D**) expression levels in the absence of arsenic stress (**A** and **B**) and following a 1 h exposure to As_2_O_3_ (**C** and **D**). NELFA-GFP is shown in green and nuclear DNA is stained with DAPI (blue). Scale bars 10 μm. See Suppl. Movie 1 for a *z*-scan view of representative cells with NELFA-GFP at high and low expressions. **E** Schematic illustration of the living-cell experiment. **F** Imaging of an unstressed HeLa cell with diffusing NELFA-GFP. Bright spots correspond to diffusing NELFA-GFP molecules imaged at different time points (as indicated). Camera exposure time 70 ms. Scale bar 10 μm. Outputs from image analysis are shown below microscopy images. NELFA-GFP clusters are shown in grey, with trailing lines identifying the track of each cluster. The number of NELFA-GFP within each cluster is shown alongside. The pixel size is 160 nm×160nm. 3 cells; 679 tracks. **G** Analogous results for a HeLa cell with NELFA-GFP upon arsenic stress. The time after arsenic exposure is indicated. After about 930s, one cluster grows irreversibly into a condensate. Camera exposure time 70 ms. Scale bar10μm. 9 cells; 3631 tracks. See Suppl. Movies 2–5 for time-resolved microscopy and image analysis of these and other cells.

Real-time tracking of cluster dynamics is not possible over extended times with Airyscan microscopes or other superresolution imaging methods because of limitations in image acquisition time, poor signal-to-noise ratios, or strong photobleaching. To image and track NELFA-GFP in living cells, we therefore used a highly inclined and laminated optical sheet (HILO) microscope [44] [Fig. 1**E**; see also Suppl. Note 3]. We used camera exposure times of 70 ms, frame rates of 0.1 s^-1^ and a low laser power to obtain sharp images and to avoid photobleaching. However, a low laser power also results in a reduced signal-to-noise ratio, rendering conventional threshold-based image analysis ineffective. We instead used a machine-learning algorithm to segment NELFA-GFP from cellular and non-cellular backgrounds, coupled with a singleparticle tracking algorithm to track individual NELFA-GFP clusters during condensate formation [Suppl. Note 4].

Using this set-up, we first imaged an unstressed living cell at intervals of 10 s. We observed the diffusion of individual NELFA-GFP spots until photobleaching occurred. Fig. 1**F** shows four images of a cell without arsenic exposure alongside the corresponding cluster tracking analysis. Full trajectories for this and two additional cells are provided as Suppl. Movie 2, and the tracking analysis for the presented cell is provided as Suppl. Movie 3.

Next, we added 100 μM As_2_O_3_ to stress the cells and trigger condensate formation of NELFA-GFP. In Fig. 1**G**, we show representative images along a trajectory for one cell as a function of time following exposure to As_2_O_3_. Many small NELFA-GFP clusters could be observed; these not only move in space, but also dynamically grow and shrink. Full trajectories for this and eight additional cells are shown in Suppl. Movie 4, and the tracking analysis for this cell is provided in Suppl. Movie 5. In all cells in which NELFA condensates formed, we found that NELFA-GFP clusters continually grow and shrink until they reach a critical size (see Suppl. Note 5 for data on all cells). However, it appears that once a cluster reaches a critical size, it continues to grow into a larger condensate, and such dynamic behaviour can therefore serve as an initial distinction between small (‘pre-condensate’) clusters and large clusters (‘condensates’). At higher expression conditions, several clusters can reach the critical size (see Fig. 1**A,B**). We investigate this nucleation-like behaviour further below.

### Fixed cells provide a calibration for living-cell data

Our microscope is capable of observing single GFPs; however, even at the lowest expression conditions studied, where only single visible condensates ultimately formed, the density within clusters soon became too high to separate and count single GFPs. We therefore combined our living-cell tracking experiments with photobleaching-step counting [45] in different fixed cells under identical expression and stress conditions, which enabled us to quantify the number of NELFA-GFP molecules in dense regions with near single-molecule accuracy. We added this quantification to the living-cell movies and images shown in Fig. 1**F,G**.

To obtain such data with near single-molecule sensitivity, we fixed HeLa cells at up to 10 different times following exposure to As_2_O_3_ and counted photobleaching steps for all clusters. This cannot readily be done in living cells, as GFPs can only be bleached once at a defined time point. Counting photobleaching steps has the advantage of counting the local number of proteins with high accuracy; by contrast, in intensitybased measurements, brightness variations in cells affect the result. Fig. 2**A–D** shows two examples of how single GFPs are counted in fixed cells over time. Since every photobleaching step corresponds to precisely one NELFA-GFP molecule, we can convert NELFA-GFP areas from living-cell imaging into the numbers of molecules in such an area, and in turn obtain the number of molecules in each cluster (see Suppl. Note 6).

**Figure 2.**
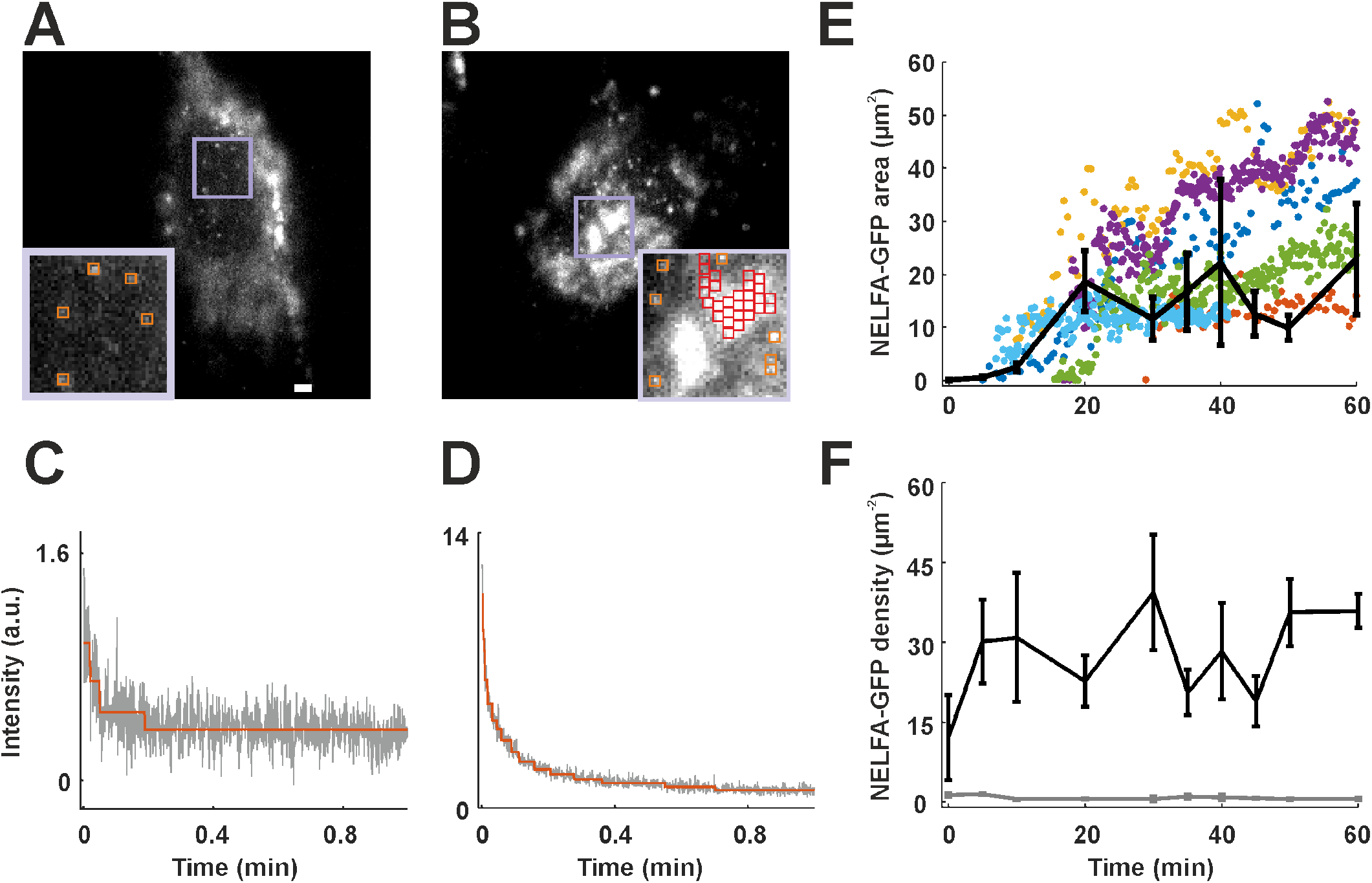
Converting areas of GFP regions into numbers of NELFA-GFP molecules. **A** Isolated GFP spots found in an unstressed and fixed HeLa cell. In the zoomed-in panel, five regions are outlined in orange. **B** Isolated and contiguous GFP regions found in a stressed and fixed HeLa cell. The condensate (i.e. the largest contiguous region) corresponds to regions outlined in red. Scale bar: 2 μm. Photobleaching steps were measured from each grid. Example curves are shown for **C** an unstressed cell and **D** a stressed cell. Steps (3 and 15, respectively) were determined by AutoStepfinder. **E** The black line gives sizes of clusters and condensates averaged over all fixed HeLa cells, while circles are tracked sizes of NELFA-GFP regions in living HeLa cells. Each colour represents a distinct cluster that ultimately grows into a NELFA-GFP condensate in living-cell experiments. **F** Density of NELFA-GFP molecules in dense (black) and dilute (grey) phases measured in fixed cells. Data from 4330 photobleaching traces and a total of 35 cells. The average number density of NELFA-GFP in the dense phase is (29±2) μm^-2^.

In Fig. 2**E**, we show that the areas of clusters at different time points for the nine living cells (coloured dots) agree well with the average data from the 35 fixed cells (black line). Such consistency is especially striking considering that the latter should be a lower limit on the condensate size, as the 35 fixed cells were selected randomly and a condensate would not have formed in all of them within 60 min. Finally, we show in Fig. 2**F** that the average density of NELFA-GFP in the dense phase (i.e. in clusters and condensates) is almost unchanging at ~29 μm^-2^, which indicates a constant saturation concentration.

In living-cell experiments, we found that a critical area of ~10 μm^2^ (see Suppl. Fig. S9) was necessary for condensate growth, which translates to ~300 NELFA-GFP (10 μm^2^ × 29 molecules μm^-2^). As only every fifth molecule was GFP-labelled (Suppl. Fig. S5), the mean critical nucleus size under these conditions was thus ~1500 NELFA molecules. The cell-to-cell variation was rather large, ranging from ~5 μm^2^ to almost ~50 μm^2^, so the critical size might be different at higher expression levels of NELFA. To validate our results, we determined the total number of NELFA molecules in the nucleus. To this end, we first determined the number of NELFA-GFP in the focal plane, which is roughly 1000 for the cell depicted in Fig. 1**F** and up to 5000 for the largest condensate observed (celll in Suppl. Movie 4). The focal plane is ~1 μm deep; for a nucleus of ~7 μm in diameter we therefore only detect about 1/5 of the molecules. Moreover, since only every fifth molecule was labelled, we multiply the number of molecules in the focal plane by 25 in total, resulting in between 25 000 and 125 000 NELFA proteins in the nucleus. This is within a small multiplicative factor of the number of NELFA in the nucleus, 155 688, determined by mass spectrometry [46], which therefore supports the quantitative nature of our analysis.

### NELF condensates are formed by non-classical nucleation

We showed above that condensate formation is a rare event and occurs only once a certain threshold size is reached [Fig. 2**E**]. To clarify the mechanism by which condensates form, we determined the lag time from the start of measurements to when fast growth to a large cluster size occurs [Suppl. Note 7]. These times are broadly distributed, which is a hallmark of a nucleation-and-growth mechanism [47]. Moreover, the larger the initial concentration of nuclear NELFA, the faster the cluster growth, as expected for a nucleation-controlled process [48]. However, the varied shapes of clusters [Fig. 1**G**] indicate that the formation of an interface between the dense and dilute phases does not result in a large disfavourable free energy, and the wide distribution of cluster sizes suggests that cluster growth is not controlled solely by a competition between a favourable bulk term and a disfavourable interfacial term.

To probe this hypothesis further, we divided cluster trajectories into those that occurred prior to the fast condensate growth (‘pre-nucleation’) and those that occurred in afterwards (‘post-nucleation’) [Suppl. Note 7]. This threshold is defined separately for each nucleus, accounting for the heterogeneity of the cells. For each scenario, we computed the probability *p_n_* that a monomer is in a cluster of a particular size ***n*** [Fig. 3**D**], and, in turn, a Landau free-energy difference between clusters of a certain size relative to monomers as Δ*G_n_* = -*k*B*T* ln(*p_n_*/*np*1). The resulting free-energy landscape [Fig. 3**E**] does initially increase with a typical surface scaling (~ *n*^2/3^), but then plateaus, suggesting the system may have a broad range of effective interaction strengths [49] and that the driving force for cluster growth is governed by a more complex mechanism than in classical nucleation theory. This plateau may in part also reflect the heterogeneity of the cellular environment; if the free energy increases for some (pre-critical) cells but decreases for post-critical ones, the overall average may appear flat, highlighting the importance of tracking and analysing individual cells. Finally, for unstressed cells, where no condensate formation has been observed, the free-energy barrier closely follows that of the stressed cells initially, but stops suddenly and does not plateau. This suggests that, although the initial cluster growth is governed by a thermodynamic disfavourability of interface formation, subsequent cluster growth is blocked in unstressed cells. We cannot determine the mechanism for this blocking at this stage, but there can be many driving forces in the complex non-equilibrium environment of the nucleus, such as a loss of valency [50] or the blocking of the DNA sequences necessary for cluster formation [51] or other biochemical interactions [38]. Below, we show that p38 kinase plays a role in this mechanism.

**Figure 3.**
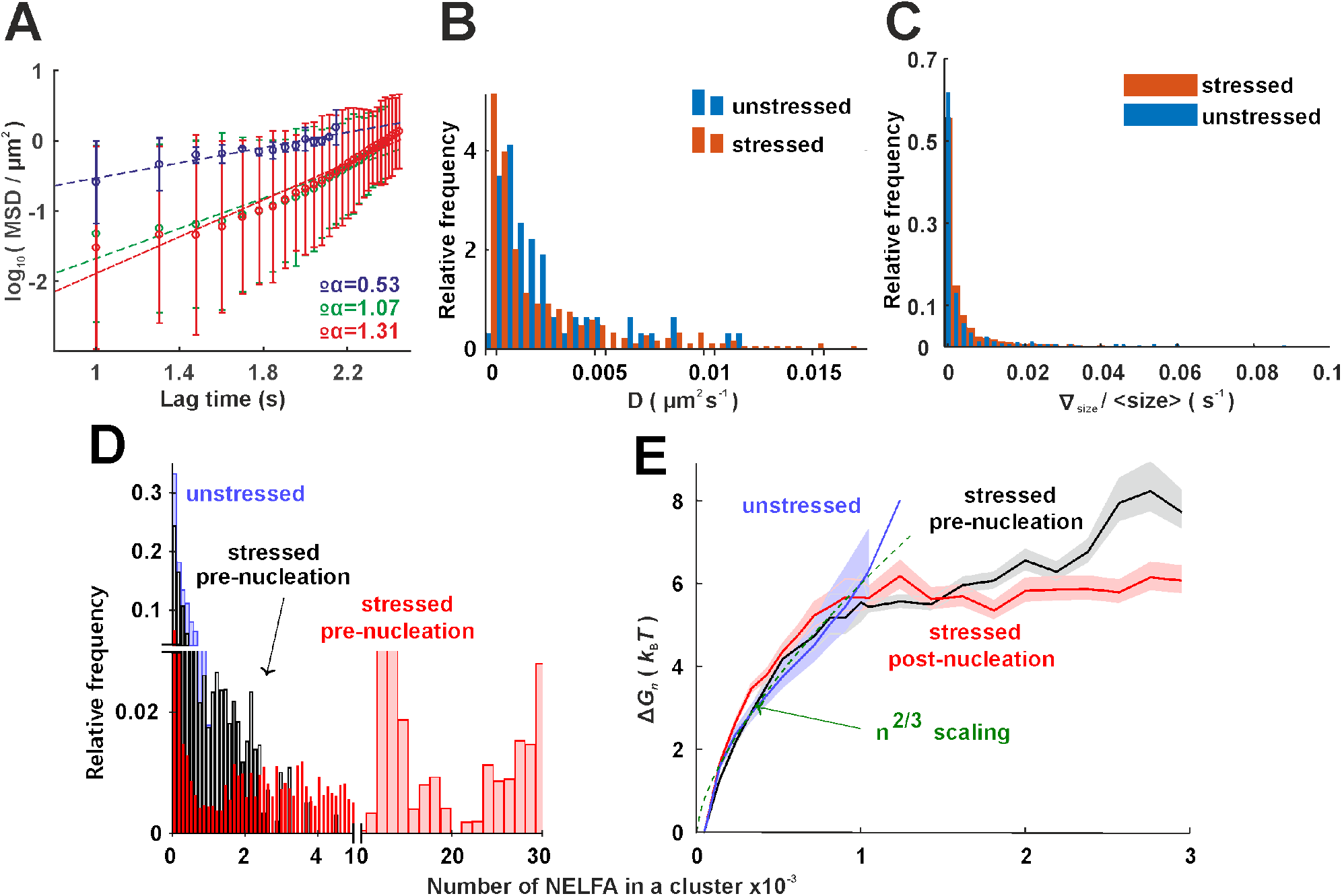
Dynamic clusters are present in stressed and unstressed cells. Large clusters are formed by non-classical nucleation. Various quantities are compared for stressed and unstressed cells: **A** The mean squared displacement (MSD); **B** the effective diffusion coefficient; **C** the scaled gradient in cluster size; **D** the probability of a monomer being in a cluster of a certain size for stressed cells prior to nucleation, after nucleation started (post-nucleation) and for unstressed cells; **E** the free energy for a certain cluster size (see main text for details). The 99 % confidence intervals shown are computed from 10 000 bootstrap samplings of data values. In **D** and **E**, 7 cells and 1262 clusters in ‘stressed’ pre- and 3028 clusters in post-nucleation, respectively; 3 cells and 679 clusters in ‘unstressed’ pre-nucleation.

Finally, we investigated further the mechanism of condensate growth post-nucleation. One possible mechanism by which large clusters could grow is Ostwald ripening [52], where small clusters gradually shrink as the largest one grows. However, we found that in our case, cluster sizes were relatively evenly distributed across a wide range of cluster sizes, and the probability of a protein being in a small cluster did not increase as one large cluster grew (Suppl. Movie 6). Moreover, we determined the mean gradient of cluster size for every tracked cluster *j* as 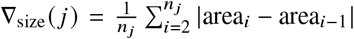, where *n_j_* is the total number of steps over which cluster *j* could be identified, and area¿ is the cluster’s area at step *i* of the trajectory. When we divided the gradient of cluster size by the area of the cluster [Fig. 3**C**], the resulting data fell within a narrow range even though the cluster size gradient itself was very much larger for large clusters than for smaller ones. This suggests that addition of proteins does not occur stepwise; instead, larger clusters sweep up more of the smaller clusters via coalescence, as opposed to Ostwald ripening (Suppl. Note 11). Similar behaviour has been observed for cortical condensates [38].

By combining fixed-cell data with living-cell imaging experiments, we can gain further insight into the real-time diffusion of all 4310 clusters investigated, for both stressed (3631) and unstressed (679) cells. Fig. 3**A** shows the mean squared displacement (MSD) of the centres of three example clusters over time from living-cell experiments. Although this graph already shows that the clusters do not only exhibit free Brownian motion, we initially calculated an effective diffusion coefficient *D* from a fit to the Einstein relation, 〈*r*^2^〉 = 4*Dt* in the long-time limit. Both stressed and unstressed cells comprise clusters with a similarly broad distribution of diffusion coefficients [Fig. 3**B**], while the background fluorescence can easily be distinguished from NELFA-GFP (see Suppl. Note 9). To quantify the extent to which clusters diffuse freely, we also used a generalized diffusion equation 〈*r*^2^〉 = 4*Dt^α^* and determined the exponent *α*. Fig. 3**A** indicates that different diffusion mechanisms were in effect, ranging from free diffusion (*α* = 1) to sub-diffusion (*α* < 1) and directed diffusion (*α* > 1). This range of behaviours is seen for both stressed and unstressed cells (Suppl. Note 10) and underlines the need to observe cluster formation in living cells, as such behaviour is not usually seen in model phase-separating systems.

### P38 kinase is required to form large clusters

The above physical analysis of cluster dynamics suggests that there might be a different regulation of small clusters compared to large clusters. To test this hypothesis, we investigated the effect of a p38 kinase inhibitor on the different clusters. P38 kinase has been shown to shuttle to the nucleus upon stress [41, 53]. To ascertain whether the p38 kinase inhibitor interferes with NELF cluster formation, we incubated living cells with the p38 inhibitor for 1 h and then added As_2_O_3_ and observed the cells for 30 min under our HILO microscope. Fig. 4**A** shows snapshots for an example cell [see Suppl. Movie 7 for this and two other cells] and Fig. 4**C** the distribution of the maximum cluster size from each tracked cluster. In the presence of the p38 kinase inhibitor, clusters larger than 600 NELFA are not formed, suggesting that p38 kinase is required for large cluster formation. In addition, this provides another way of distinguishing ‘small’ and ‘large’ clusters: we previously showed how cluster dynamics can be used as a criterion, but one also could use the susceptibility towards the p38 kinase inhibitor.

**Figure 4.**
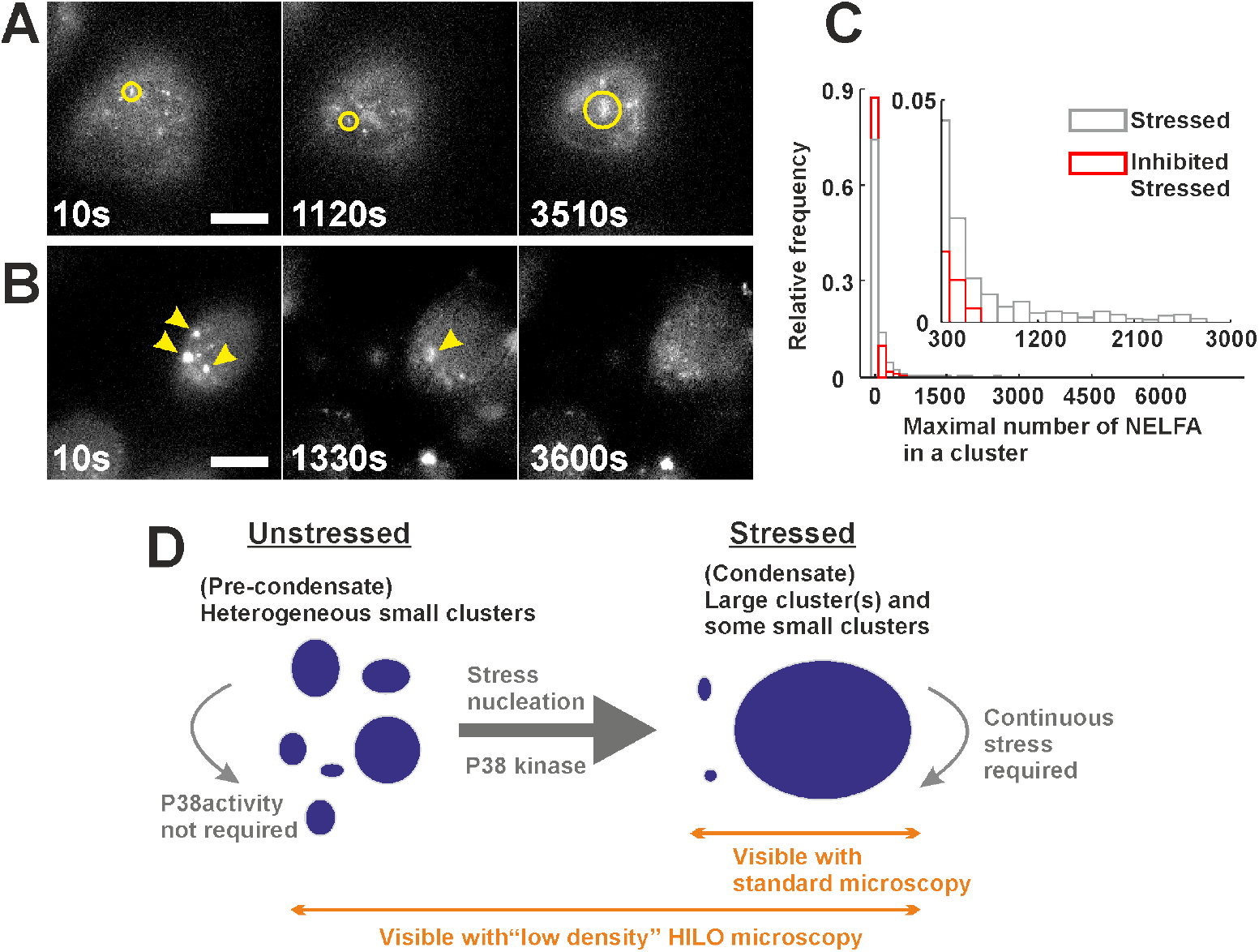
P38 kinase is required to form large clusters. **A** Cell nucleus of a cell that was treated with a p38 kinase inhibitor for 1 h prior to the addition of arsenic. Imaging with the HILO microscope began when As_2_O_3_ was added. Small clusters assembled, but no large clusters were observed, even after 1 h of arsenic treatment. 3 cells. **B** Cells were exposed to As_2_O_3_ for 30 min to form condensates (yellow arrows). The medium was exchanged (i.e. arsenic removed) to include p38 inhibitor at time 10 s, which caused several condensates (large clusters) to disassemble, while small clusters were still present. Images were corrected for photobleaching effects using the ImageJ plugin (Histogram Matching). A total of 13 cells were investigated; about half of them showed disassembly [see Suppl. Movie 8 for an example]. Disassembly of already formed large clusters was never observed in the presence of arsenic. All scale bars 10 μm. **C** Analysis of the maximum cluster sizes reached by each tracked cluster with and without pre-incubation with a p38 kinase inhibitor upon stress. The inhibitor interferes with the formation of large clusters, as no clusters larger than 600 NELFA were observed, providing another distinction between ‘small’ and ‘large’ clusters in living cells. 3 cells; 344 clusters. **D** Schematic illustration of how small and large clusters respond to stress and how p38 kinase interferes with this process (see main text for details).

Finally, we tested what is required for the maintenance of large clusters. To this end, we first exposed cells to As_2_O_3_ for 30 min to form condensates and then added p38 inhibitor in the presence of arsenic, i.e. the cells were still under stress. None of the condensates disappeared (4 cells), but half the cells died. We next exposed cells to As_2_O_3_ for 30 min to form condensates, but now we removed the stress by exchanging the medium, and added the p38 inhibitor; this led to several large clusters dissolving (Fig. 4**B**). The cell-to-cell variation for this effect is high: 7 of the 13 investigated cells show full or partial disappearance of large clusters, but some also stay intact (see Suppl. Movie 8 for an example). By contrast, we never observed the dissolution of large clusters in the presence of arsenic. Altogether, our data show that p38 kinase is required to form large clusters under stress, and thus seems to be involved in unblocking the nucleation of NELF clusters in stressed cells. On the other hand, once large clusters have formed, they remain present for as long as the stress environment is kept, independent of the p38 kinase.

## CONCLUSION

A combination of fluorescence imaging in living and fixed cells allowed us to quantify the dynamics of cluster and condensate formation for NELF in living cells. We obtained time-resolved NELF cluster sizes with almost single protein resolution and could follow the clusters’ dynamics in real time. We showed that large stable clusters (condensates) only formed rapidly in the nucleus of stressed cells once a threshold of ~1500 NELFA proteins was reached, albeit with considerable variability amongst different cells. Strikingly, before condensate formation and even under subsaturation conditions where no condensates form, many smaller clusters of tens to hundreds of NELFA proteins were observed.

Our physical analysis of all clusters in living cells shows that classical nucleation theory is insufficient to describe NELFA nucleation. We obtained a relatively flat free-energy landscape following an initial surface-dominated barrier, with the probabilities of molecules being in clusters of many different sizes surprisingly similar. It appears that nucleation in unstressed cells is likely not blocked by a high free-energy barrier, but by another mechanism, for example interactions with other proteins. P38 kinase could be one such protein, as we have shown that it is involved in the release of the blocking of the formation of large NELF cluster in the nucleus. As a p38 inhibitor had no measurable effect on the small NELF cluster, the response to it provides another means of distinguishing between small and large clusters. Future experiments of the type we have presented will reveal if these transient small clusters are also regulated by chaperones, which have already been shown to modulate size distributions of self-associating proteins [54], even in an ATP-dependent manner [55].

In summary, we have shown that small, transient, dynamic clusters of NELFA appear before stable condensates form, even in unstressed cells. These small clusters differ from the large ones in their dynamics and in their response to the p38 kinase inhibitor. We expect that small dynamic clusters and the blocking of nucleation play an important role in cellular regulation and signalling. Such blocking may for example allow for a significant build-up of mass that can then result in rapid condensation when the cell requires it.

## MATERIALS AND METHODS

### Cell culture and induction of NELFA-GFP expression

NELFA-GFP stable HeLa were grown in DMEM (Gibco 31053-028) to which 10% FBS (Gibco 10270-106), 100 units/mL penicillin (Sigma P4333), 100 μg mL^-1^ streptomycin (Sigma P4333) and 2mM L-glutamate (Sigma G7513) were added, at 37 °C and 5 % CO_2_ [42]. Between 1×10^4^ and 2×10^4^ cells were seeded in Ibidi dishes (μ-dish 35 mm, high glass bottom dishes, Ibidi, 81158) with a refractive index of 1.52 to grow for 24 h to 30 h before tetracycline induction. NELFA-GFP expression was induced by tetracycline (0.2μg mL^-1^, 0.4μg mL^-1^ and 1 μg mL^-1^) for 4h to 6h before living-cell imaging or fixation.

### HILO microscopy and living-cell imaging

Cells were imaged while they were exposed to As_2_O_3_ (within 75 min) in living-cell imaging solution (Invitrogen, A14291J) at 37 °C on a custom-built fluorescence microscope in HILO mode (objective: Nikon Apo TIRF 100×/1.49 oil) with an EMCCD camera (Andor iXon Ultra 897) at a laser (Coherent, 473 nm) excitation power density of 60 mW cm^-2^ (for time-lapse living-cell imaging) or constant 240 mW cm^-2^ (for measuring the photobleaching steps in fixed cells). The recorded area was 40.96 μm×40.96 μm; see also Suppl. Note 3.

For our single cell experiments we selected HeLa cells in which NELFA-GFP fluorescent spots diffused in an apparently random manner. Controls have shown that the background fluorescent pattern was either static or moved in a directed manner. We immediately added 100 mM As_2_O_3_ (Sigma 202673-5G) to the selected living-cell imaging solution at multiple positions of the Ibidi dish, which contained 2 mL living-cell imaging solution, resulting in a final concentration of 100 μM of As_2_O_3_. The time lapse was started when As_2_O_3_ was added, with a time interval of either 10 s or 30 s between two frames. The camera exposure time of each frame was 70 ms.

For fixed-cell imaging for measuring photobleaching steps, HeLa cells were treated with 100 μM As_2_O_3_ for 60 min and were further fixed using Image-iT™ fixative solution (Invitrogen FB002) at room temperature for 15 min. Cells were then washed with DPBS (Gibco 14190144) five times, after which they were ready for imaging. The fixed cells for analysis were selected to have similar NELFA-GFP regions compared to the living cells at the respective time points.

Finally, we selected cells which had ideal expression conditions of NELFA-GFP for our fluorescence experiments, i.e. about one NELFA-GFP per four NELFA. We believe that this minimally perturbs the wild-type system, which is supported by our viability assays. In addition, every single cell from our living-cell experiments was observed for about 60 min. Therefore, the full set of results (dynamic size, diffusion coefficients) was obtained for every single cell, without averaging. We do not claim that every single cell shows exactly this dynamic behaviour, but many cells do, and in total we have investigated more than 60 cells.

To exclude imaging artefacts, we measured a control system (2NT-DDX4-GFP) [56] with our setup and find spherical condensates (Suppl. Fig. S7**C**).

### Analysis of living-cell imaging movies

Recorded movies were first processed using the Weka segmentation plugin in Fiji [57] to extract NELFA-GFP regions from the manually assigned cellular and non-cellular background and were further processed using the Mosaic plugin [58, 59] in Fiji to track NELFA-GFP regions (Suppl. Note 2). Further data were evaluated and plotted using the ImageJ macro (open source) and MATLAB (MathWorks) using custom code.

### Analysis of fixed-cell imaging movies

Recorded movies were first analysed using Fiji, ImageJ macro and MATLAB custom code to extract locations and bleaching curves in dilute and dense phases with background correction (Suppl. Note 6). Photobleaching steps were measured using AutoStepfinder [60].

### Immunofluorescence microscopy

NELFA-GFP HeLa cells seeded on Ibidi chambers were treated with tetracycline for 4 h to induce the expression of NELFA-GFP. Some cells were further exposed to 100 μM As_2_O_3_, as described previously. Cells were washed with PBS and fixed in 4 % paraformaldehyde in PBS for 10 min at room temperature, followed by permeabilization with 0.3 % Triton X-100 in PBS for 10 min at room temperature. Cells were washed with PBS and incubated with DAPI (Sigma, #D9542; 1: 1000 from 0.5 mg mL^-1^ stock) for 5 min at room temperature. Thereafter, cells were washed with PBS and stored at 4 °C before imaging. Fluorescence images were generated using a Zeiss LSM800 microscope equipped with a 63×, 1.4 NA oil objective and an Airyscan detector and processed with Zen blue software and ImageJ/Fiji. Cells were imaged as z-stack with 130 nm sections with a lateral resolution of 120 nm.

## Acknowledgements

We thank Ibrahim Cissé, Rosana Collepardo-Guevara, Stephan Grill and Rohit Pappu for helpful discussions, and Adam Klosin (Hyman lab) for supplying us with the plasmid of DDX4.

This work was supported by the European Research Council (grant agreement No. 681891) and the Deutsche Forschungsgemeinschaft (DFG) under Germany’s Excellence Strategy (CIBSS EXC-2189 Project ID 390939984) and the SFB1381 programme (Project ID 403222702).

## Author contributions

A.R., R.S., T.H. designed the research; C.L., J.K., F.A., S.U. performed the measurements; C.L., A.R., R.S., T.H. analysed the data after consultation with R.G. and A.A.; C.L., A.R., R.S., T. H. wrote the manuscript. All authors discussed the results and commented on the manuscript.

## SUPPLEMENTARY INFORMATION

### Supplementary movies

**(see https://zenodo.org/record/6946008#.Yuaj7y8etdg)**

Movie 1 Z-scan view of Fig. 1. The nuclear lamina is stained by AF647 and appears in red. Scale bars 10 μm.

Movie 2 Three unstressed cells with NELFA-GFP. Scale bars 10 μm.

Movie 3 Machine-learning-based tracking of NELFA-GFP regions (without arsenic exposure).

Movie 4 Nine cells showing NELFA-GFP phase separation at low expression levels. Scale bars 10 μm. Orange rectangles mark regions of interest for the tracking of NELFA-GFP regions.

Movie 5 Machine-learning-based tracking of NELFA-GFP regions (with arsenic exposure).

Movie 6 Time-resolved cluster size probability distribution in stressed and unstressed cells.

Movie 7 Cluster formation in the presence of p38 inhibitor. Scale bars 10 μm.

Movie 8 Dissolution of condensates when stress is released (the medium is exchanged) and p38 inhibitor is added. Scale bars 10 μm.

### Note 1: Measurement of the ratio of NELFA-GFP to endogenous NELFA

HeLa cells were cultured in six-well plates as mentioned in the Materials and Methods, and NELFA-GFP expression was induced by tetracycline (1 μg mL^-1^) for four and six hours. Cells were harvested and washed in PBS. Cell pellets were frozen in liquid nitrogen and stored at −80 °C until Western-blot analysis. Cell pellets were thawed gradually on ice and lysed in lysis buffer (50 mM Tris-HCl pH 6.8, 2 % SDS and 10 % glycerol). Cellular lysates were analysed by SDS-PAGE followed by immunoblotting using antibody for NELFA (1: 500; Santa Cruz sc-23599). Signals were quantified by densitometry using Image Lab (Bio-Rad Laboratories).

**Figure S5.**
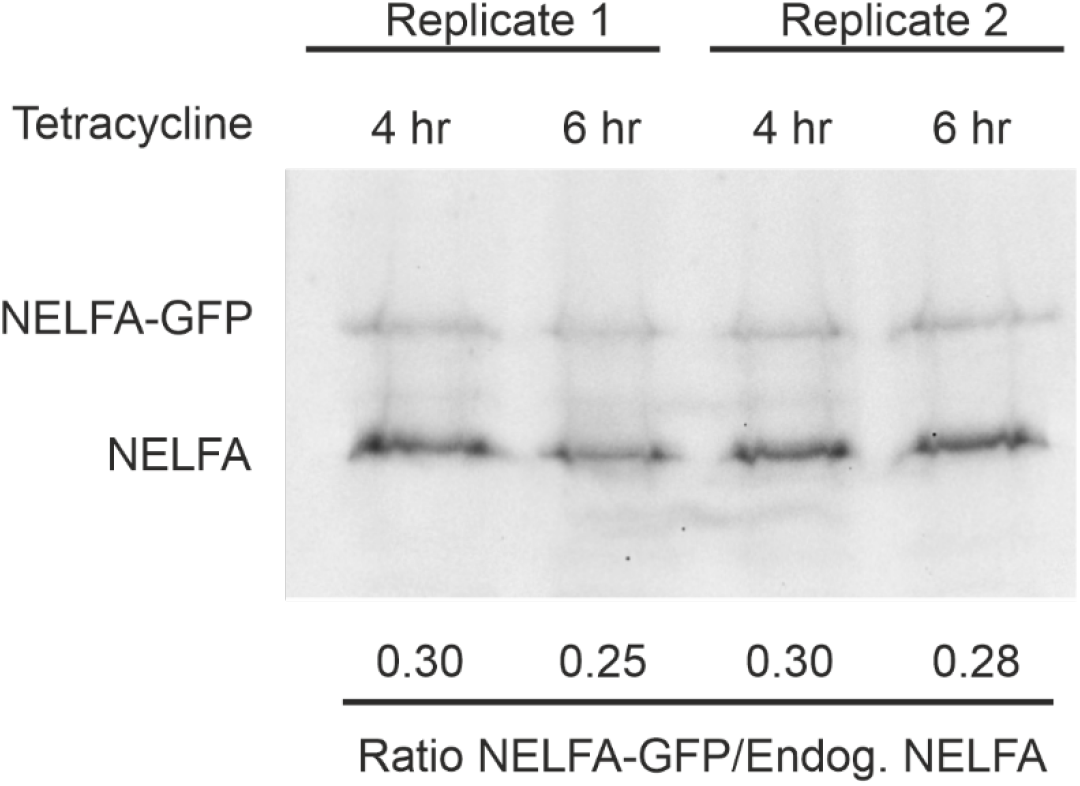
Western-blot analysis to measure the ratio of NELFA-GFP to endogenous NELFA. NELFA-GFP expression was induced with tetracycline for 4h to 6 h. Two replicates were run in the same gel. Endogenous NELFA serves as a protein loading control.

### Note 2: Measurement of the viability of HeLa-NELFA by trypan blue assay

**Figure S6.**
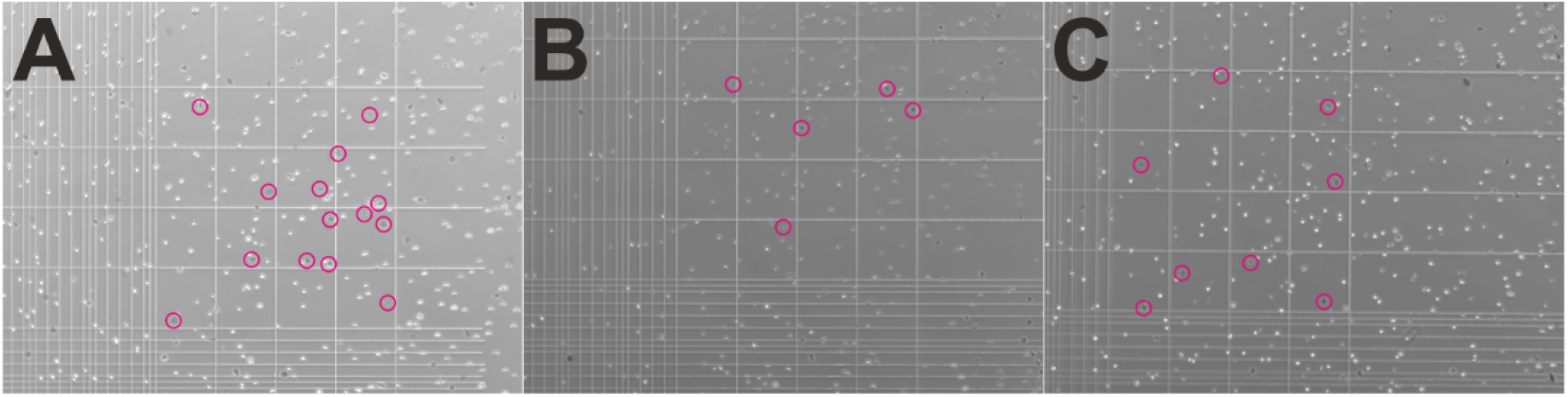
Representative bright-field images of fresh and As_2_O_3_-treated HeLa-NELFA cells stained by trypan blue. Living and adherent HeLa-NELFA cells were washed by DPBS, detached by trypsin, centrifuged (100g, 5 min, 4 °C), and resuspended in 1 mL DPBS. 100 μL suspended cells were treated by 100 μL trypan blue for 3 min. Bright (living) and dark (dead, marked by circles in magenta) cells were imaged and counted in a haemocytometer. **A** The viability ratio in fresh cells was 86%. **B** The viability ratio in cells following a 1 h As_2_O_3_ treatment of cells growing in Invitrogen living-cell imaging solution was 95%. **C** After As_2_O_3_ treatment, cells were washed by DPBS three times and grown in cell culture media for 24 h. The viability of cells after this treatment was 94%.

### Note 3: Setup and living-cell imaging of synthetic NELFA-GFP at high expression levels (24 h for NELFA-GFP expression) and synthetic 2NT-DDX4-GFP

**Figure S7.**
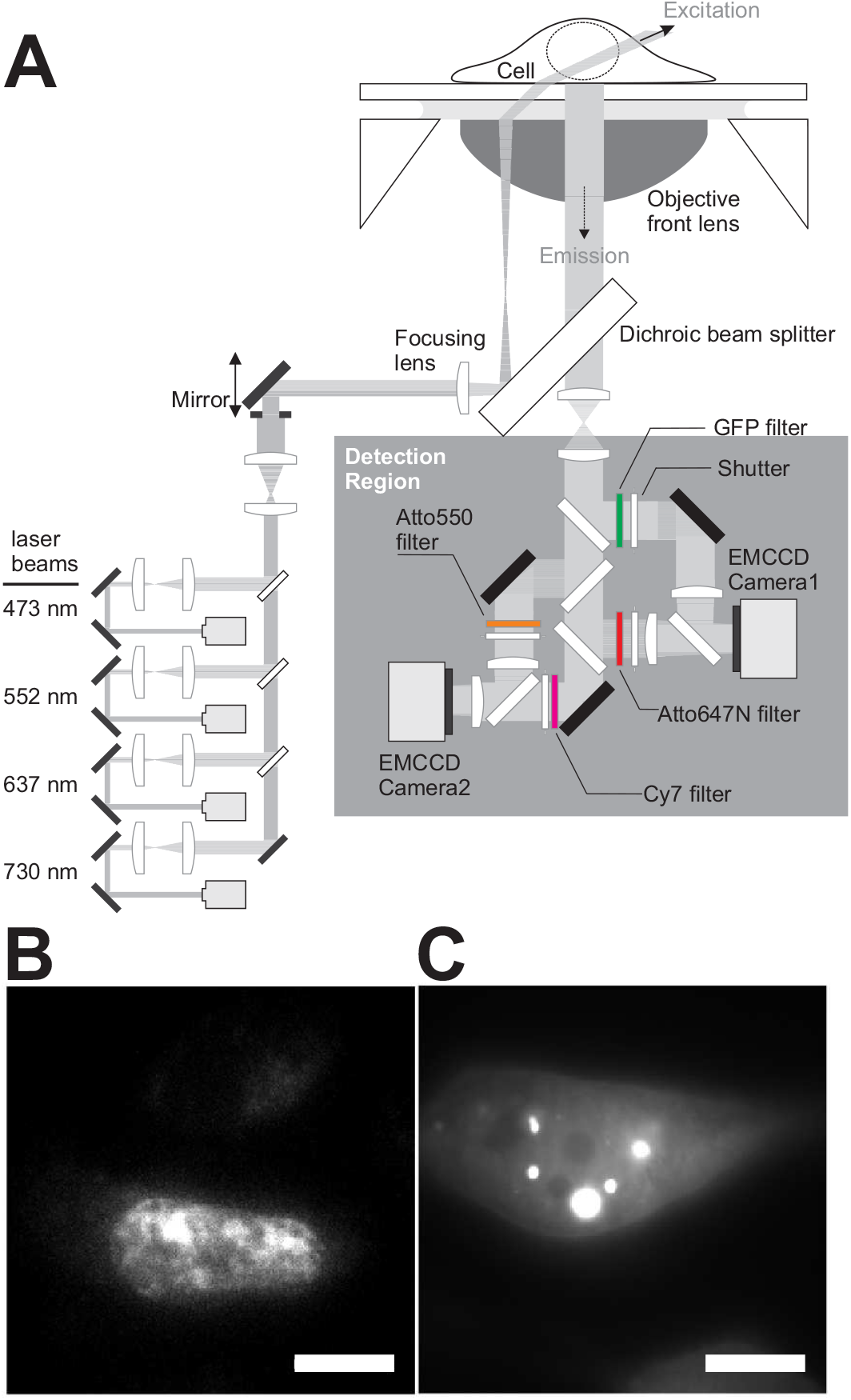
**A** Schematic illustration of the custom-built TIRF/HILO microscope for single-molecule imaging or biological phase separation in living cells. **B** A living HeLa cell expressing the NELFA-GFP (24 h of expression time) followed by 1 h of arsenic exposure. **C** A living HeLa cell expressing synthetic 2NT-DDX4-GFP (6h after transfection) [56] undergoing phase separation using the TIRF/HILO microscope. Scale bar 10 μm.

We usually only observe the formation of a single (final) condensate. At first, this may appear to contradict previous observations of several condensates for NELFA [42] as well as other publications on different proteins. In order to make sure that we do not miss other condensates in the cell, in other imaging planes, we recorded 3D Airyscan movies and confirmed that there was indeed only one large condensate under the low NELFA-GFP expression conditions used here. The formation of a single condensate is consistent with a nucleation-and-growth model requiring a critical nucleus size of around 1500 NELFA molecules. As the first post-critical nucleus grows quickly, it depletes the reservoir of NELFA molecules in the system, leaving insufficient molecules to form a second post-critical nucleus. By contrast, at the expression conditions used in our previous publication [42], there are several times more proteins in the cell, which suffices to form several condensates.

### Note 4: Machine-learning-based tracking of irregular shapes of NELFA-GFP regions

In our living-cell imaging results, images in movies had a low signal-to-noise ratio as a trade-off to avoid significant photobleaching of GFPs. To segment regions of NELFA-GFP better, we resorted to a combination of machine-learning-based segmentation algorithms, trainable Weka segmentation (Weka), a Fiji plugin [57]. First, we set three training features (Gaussian blue, Hessian and Membrane projections). Next, as illustrated in Fig. 1, we set three classifiers to determine regions of NELFA-GFP, regions of cellular background, and regions of non-cellular background. By manually delineating borders of each type of region in several frames, we trained the Weka until the machine-learning-based recognition of regions was at the level of eye detection. The output of Weka was a movie comprising binary images for each type of region.

**Figure S8.**
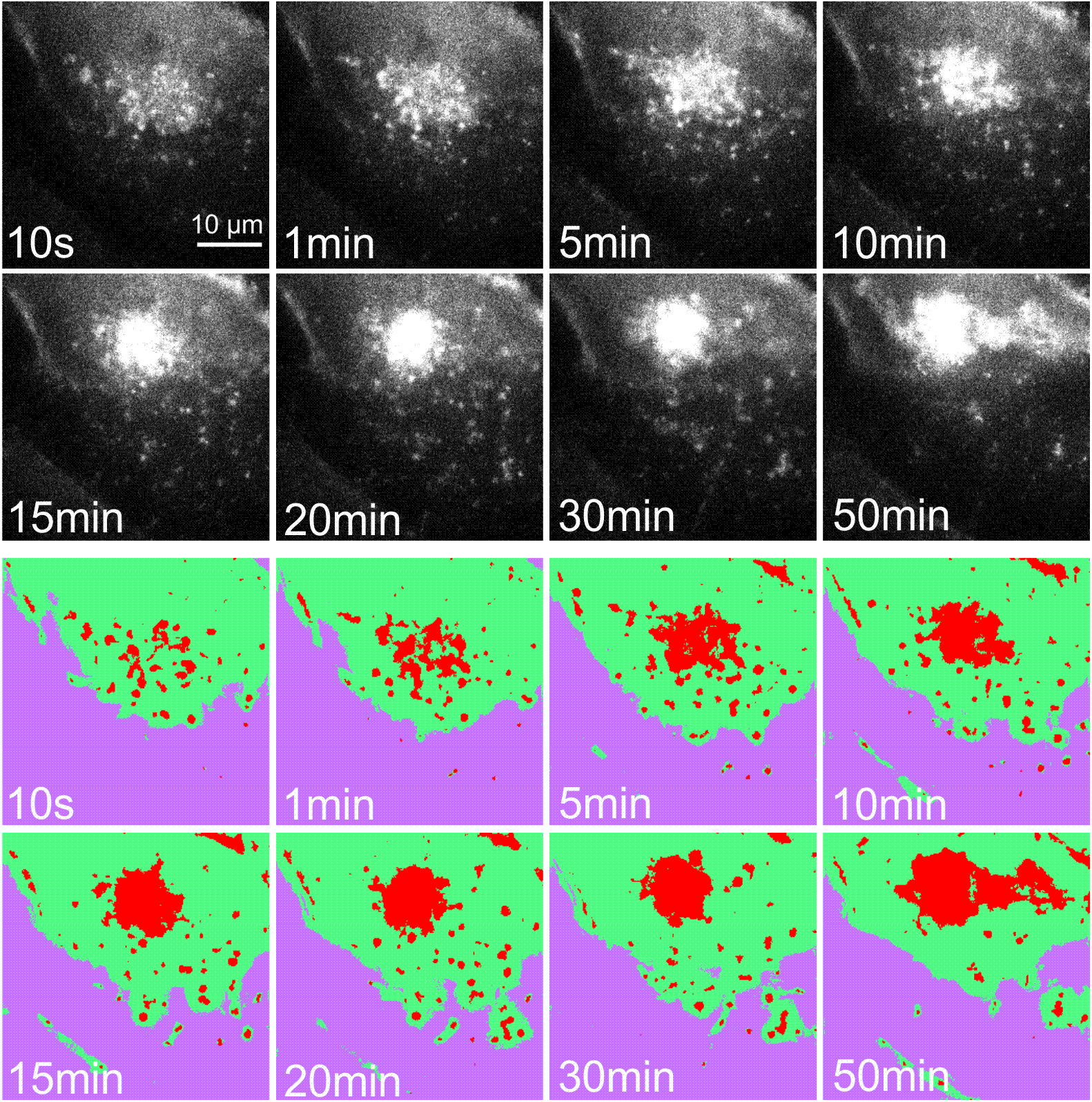
Machine-learning-based segmentation of images with trainable Weka segmentation. In eight images following the training by Weka, red regions are NELFA-GFP, green regions are regions of cellular background, and purple regions are regions of non-cellular background, respectively.

Once NELFA-GFP regions from Weka were obtained without backgrounds, we further used MOSAICsuite, another plugin in Fiji, to track NELFA-GFP regions automatically, accounting for the constantly changing region shapes. To enable reliable tracking, we first processed the binary images of NELFA-GFP regions in the sub-plugin Squassh in MOSAICsuite to perform a post-segmentation for the subsequent MOSAIC tracking [58, 59]. We used the following setting in Squassh:

1. in Background Subtraction, disable ‘Remove Background’;
2. Regularization (>0) ch1 & ch2 = 0.2 and Minimum Object Intensity Channel 1 & 2 = 0.3 for most movies;
3. Select the ‘Exclude Z edge’ option;
4. Select ‘Automatic’ in Local Intensity Estimation and ‘Poisson’ in Noise Model;
5. Values are all set to be 1 in the PSF Model;
6. ‘Remove Region with Intensities’ should be less than zero and ‘Remove Region with Size’ should be less than 2.

The Particle Tracker 2D/3D further processed the output of Squassh in the MOSAICSuite with Link Range set to 2 and Displacement set to 20, respectively [59].

### Note 5: Representative data sets of cluster sizes over time in living stressed and unstressed cells

**Figure S9.**
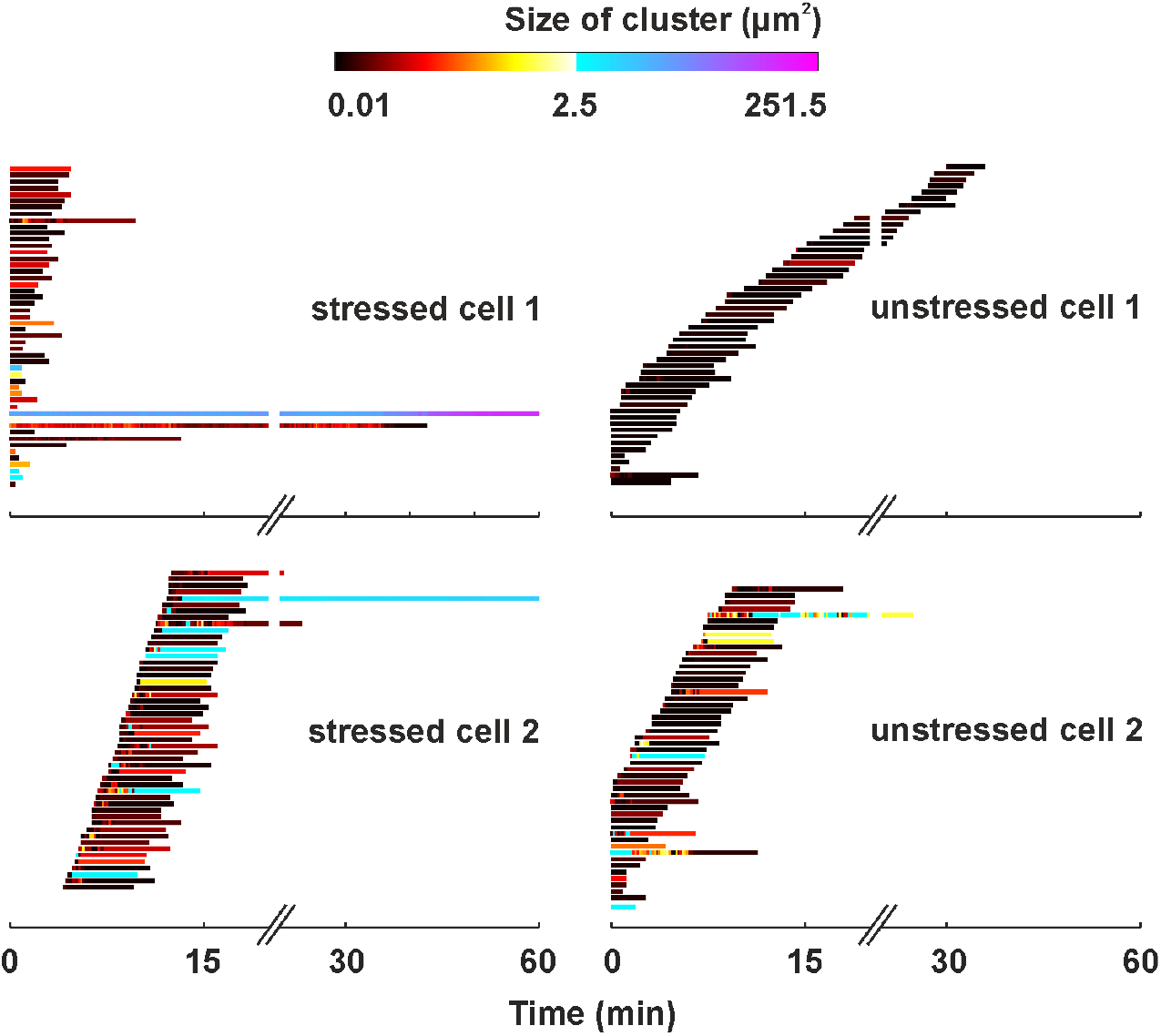
Representative changes in sizes of clusters over time in living cells.

### Note 6: Automated photobleaching-step counting in fixed HeLa cells

To count the number of NELF-GFP molecules, we measured photobleaching steps from nuclear GFP regions by imaging HeLa cells over time (200 s or 400 s) and photobleached GFP (with a 473 nm laser power of 2 mW measured at the laser set).

In a movie, GFP regions first appeared as either isolated spots or contiguous regions (Fig. 2**A-B**). To extract stepwise photobleaching curves, we first set a threshold, such that most isolated spots appeared with a size of 3 × 3 pixels in the first frame of the movie. This is because the theoretical diameter of an Airy disk of one GFP spot is 1.22 × 509 nm|1.49 = 417 nm, with the fluorescence being at ~509 nm and the numerical aperture of the objective being 1.49. This is about three pixels in one image (one pixel = 160nm×160nm).

To account for the background, we subtracted the average of the surrounding pixels from the 3 × 3 area, i.e. we assigned square grids of 5 × 5 pixels containing 3 × 3 pixel regions. The final fluorescence intensity *I*_net_ of a 3 × 3 grid in each frame was then calculated by subtracting the intensity of the region immediately surrounding the 3 × 3 region, as well as the non-specific background from the cell, namely

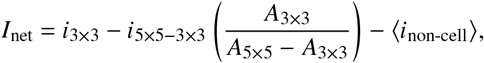

where *i* is the mean grey value in a grid of 3 × 3 pixels (*i*_3×3_) or of the 16 remaining pixels in the 5 × 5 grid surrounding the 3 × 3 region (*i*_5×5-3×3_) pixels, as appropriate, *A* is the area of the grid and 〈*z*_non-cell_〉 is the mean grey value from 10 randomly selected grids (with sizes between 3 × 3 pixels and 5 × 5 pixels) outside the cell [61].

For each photobleaching trace (without filtering), photobleaching steps were determined by AutoStepfinder (Fig. 2) [60]. Steps from AutoStepfinder should be consistent with those determined by hand by observing multiple simulated traces with various steps. The value of photobleaching steps is the maximal number of NELF-GFP molecules in each grid.

We define the ‘dilute phase’ of NELF as the region where only isolated spots with a size of 3 × 3 pixels were present, and the ‘dense phase’ as the largest contiguous GFP region with a size larger than 3 × 3 pixels). In other words, this is the largest phase-separated domain within a HeLa nucleus.

In the dense phases, the number density of NELF-GFP molecules can be calculated as the total number of molecules (determined by the total number of photobleaching steps) divided by the total area of the grids, *A*_cluster_ = *N*_grids_ × 0.23 μm^2^. For dilute phases, the density of NELF-GFP molecules is the total number of steps for all dilute-phase regions divided by the total area of the nuclear region 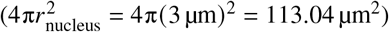, with the area of the dense phase subtracted [56].

### Note 7: Analysis of the onset of NELFA-GFP condensation in living-cell imaging movies

**Figure S10.**
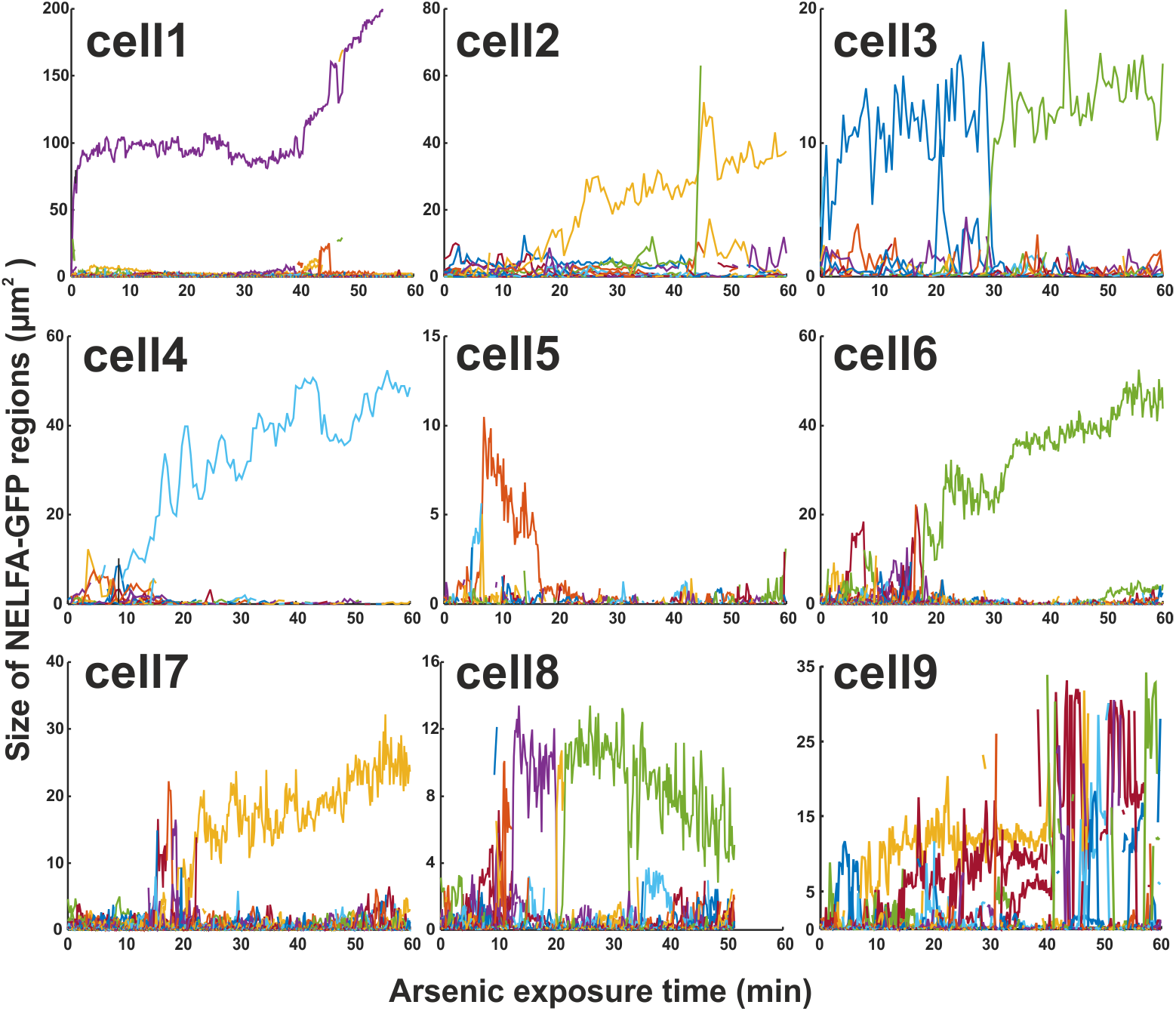
Changes in cluster sizes after arsenic exposure from the regions of interest in cells shown in Suppl. Movie 4. In total, 3631 clusters were tracked. For Fig. 3**D** and Fig. 3**E**, the choices of threshold times to delineate pre- and post-nucleation behaviour for each cell are 0.4 min (cell1), 21 min (cell2), 30 min (cell3), 13 min (cell4), 22 min (cell6), 21 min (cell7) and 7 min (cell9), respectively.

### Note 8: Diffusion coefficients and gradients in cluster size

**Figure S11.**
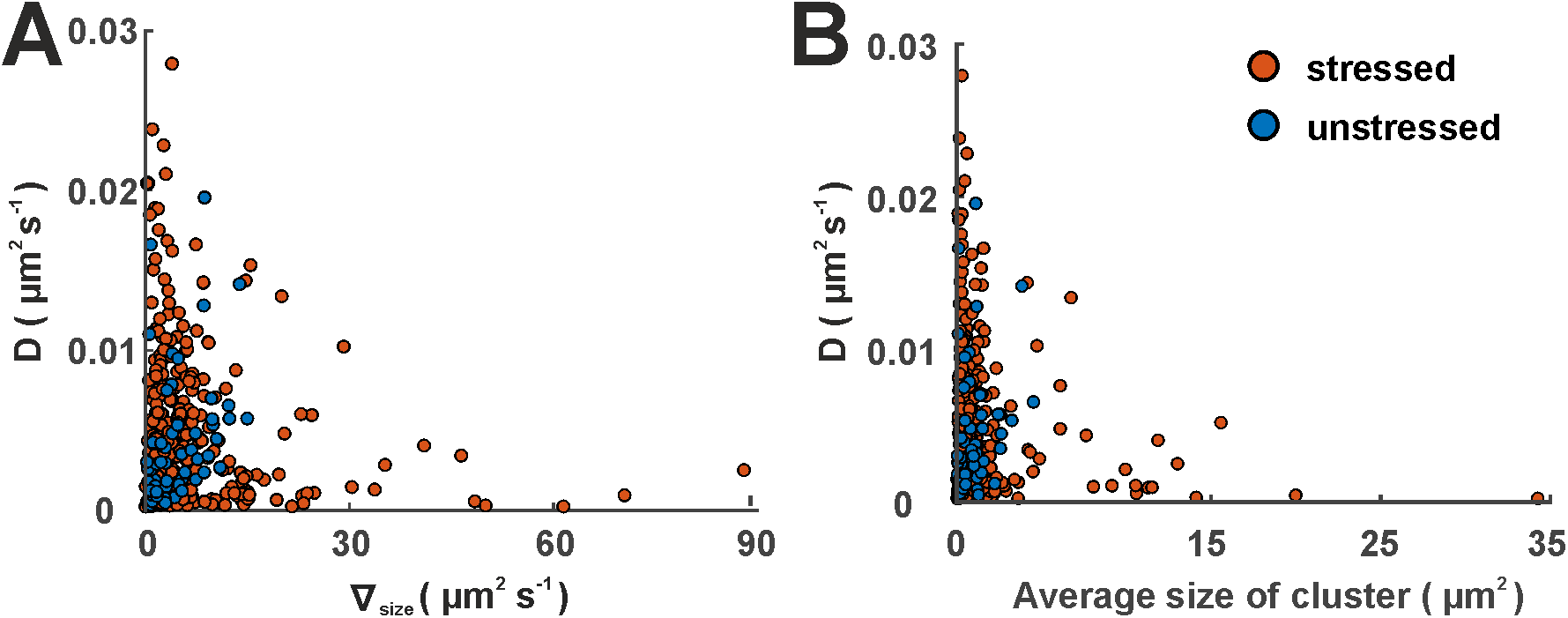
The gradients of size (**A**) and the average sizes of clusters (**B**), with associated effective diffusion coefficients.

We show in Fig. S11**A** the effective diffusion coefficient *D*(*j*) (computed by a fit to the Einstein relation, 〈*r*^2^〉 = 4*Dt* [62]) and size gradient **∇**_size_ (*j*) for every cluster *j*; the two results are only very weakly correlated, if at all, with Pearson coefficients of −0.09 (−0.15 to −0.02 at 95 % confidence) for stressed and 0.35 (0.05 to 0.57 at 95 % confidence) for unstressed cells. The change in cluster size due to possible diffusion in and out of the imaged volume is therefore negligible. Although some small part of the apparent change in size might thus have been caused by diffusion, this would not appreciably affect our results.

There are more clusters with smaller diffusion coefficients in the stressed cells, consistent with the higher probability for larger clusters; however, the diffusion coefficient is not strictly related to cluster size [Fig. S11**B**].

### Note 9: Diffusion coefficient for background compared to NELFA-GFP

**Figure S12.**
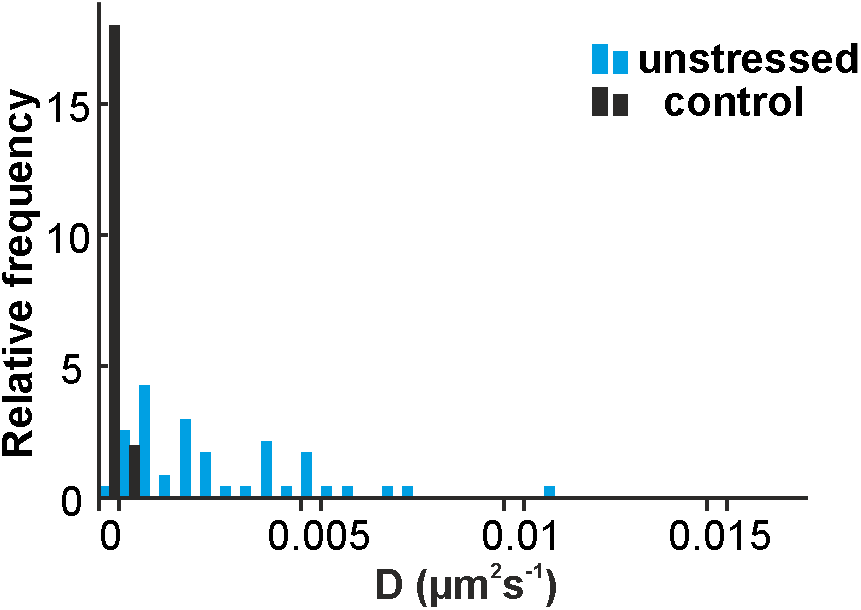
Diffusion coefficient for background compared to NELFA-GFP. Diffusion coefficients are measured from 679 tracks of three control (wild-type) HeLa cells and from 184 tracks of three unstressed HeLa cells, respectively

### Note 10: Distribution of diffusion exponents

**Figure S13.**
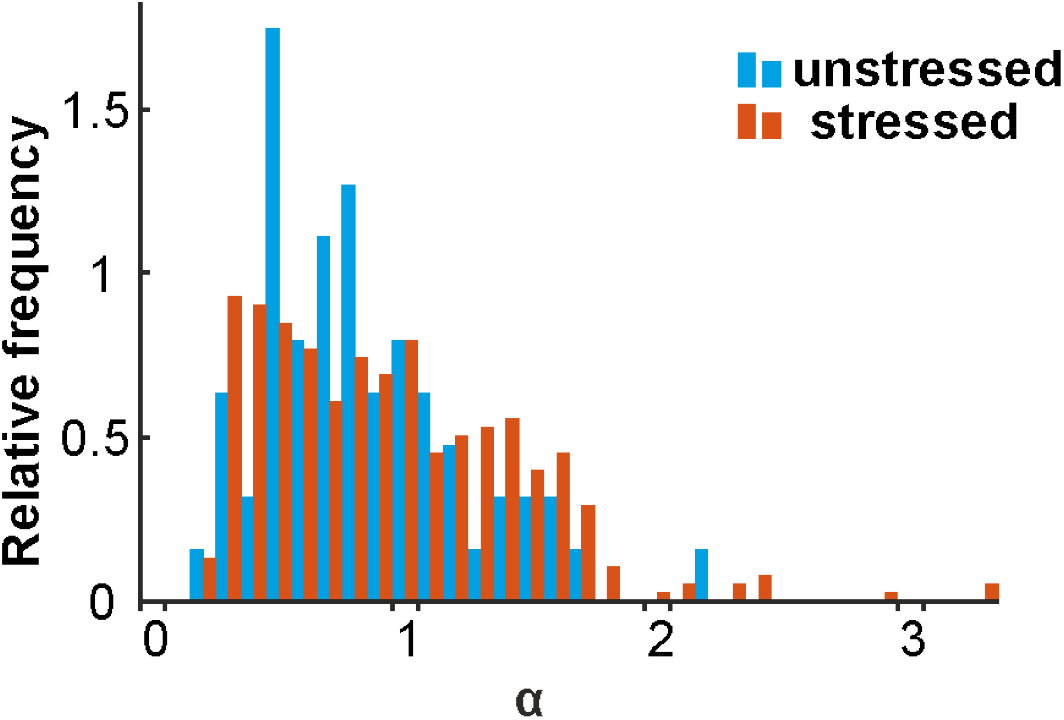
Diffusion exponent *α* for clusters in stressed and unstressed cells.

### Note 11: Coalescence of clusters

**Figure S14.**
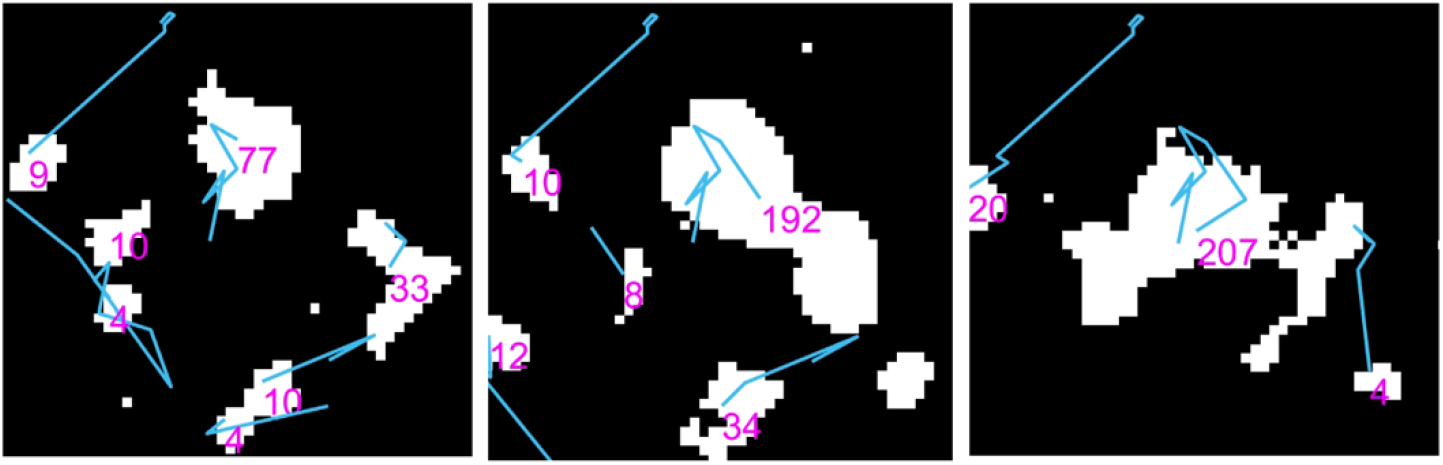
Possible coalescence of clusters. Taken from Suppl. Movie 5 at times of 120 s, 130 s and 140s.

